# Rank Normalization Enables Cross Pathway Comparison in Degradation Resolved Proteome Turnover Analysis

**DOI:** 10.64898/2026.09.30.755414

**Authors:** Zi Gao, Qiang Xiao, Jeffery W. Kelly, Evan T. Powers

## Abstract

Protein homeostasis, or proteostasis, is essential for cellular health, and its disruption contributes to cancer, aging, and neurodegenerative disease. Proteostasis is maintained largely by two intracellular degradation systems: the ubiquitin proteasome system (UPS) and the autophagy lysosomal pathway (ALP). Here, we combine pulse stable isotope labeling by amino acids in cell culture (pSILAC) with selective inhibition of the UPS or ALP to quantify pathway-dependent protein turnover and identify substrates that are preferentially cleared through each route. Direct comparison of these perturbations is complicated by their markedly different response distributions. To address the mismatch in dynamic range, we developed a normalization strategy based on within-condition ranks. Protein stabilization values are converted to percentile scores before comparison across pathways, placing both perturbations on a common relative response scale. Across two independent pSILAC experiments, 5,518 proteins were quantified in at least one condition, and 3,814 yielded high confidence half-life measurements in control, proteasome-inhibited, and lysosome-inhibited conditions. Rank normalization resolved pathway sensitivities that were obscured by direct comparison of absolute half-life changes. Importantly, the method reports relative pathway preference rather than absolute pathway contribution, exchanging effect size magnitude for comparability across perturbations with unequal response distributions. Integration with a curated proteostasis network annotation revealed coherent pathway preferences, including coordinated stabilization of proteasome core and regulatory subunits following lysosomal inhibition. This finding provides proteome-wide quantitative evidence for proteaphagy that complements previous targeted observations. Together, these results establish rank normalization as a practical strategy for comparing degradation resolved proteome turnover across perturbations with different dynamic ranges.

## Introduction

Protein homeostasis, or proteostasis, encompasses the processes that control protein abundance, conformation, interactions, and localization across the proteome. These processes are coordinated by the proteostasis network, which balances protein synthesis, folding, trafficking, and degradation throughout the protein life cycle. Disruption of this network contributes to a broad spectrum of diseases, including cancer, aging related disorders, and neurodegeneration^1–3^.

At its core, proteostasis depends on the continuous balance between protein synthesis and degradation. Two proteolytic systems dominate intracellular protein turnover in mammalian cells. The ubiquitin-proteasome system (UPS)^4^ preferentially removes many short lived, misfolded, and regulatory proteins after ubiquitin conjugation. The autophagy-lysosome pathway (ALP) mediates both bulk and selective degradation of proteins, macromolecular complexes, and organelles^5, 6^, though lysosomes also degrade other classes of biomolecules. Determining the degradation route used by individual proteins is therefore fundamental to understanding proteome maintenance, but it remains analytically challenging.

Conventional measurements of steady state protein abundance cannot readily distinguish whether a change after pathway inhibition reflects altered synthesis, altered degradation, or both. Pulse SILAC addresses this limitation by resolving newly synthesized and preexisting protein pools over time. Following a switch from light to heavy amino acids, decay of the preexisting light pool provides a kinetic measure of protein loss and enables estimation of protein specific degradation rates and half-lives^7, 8^. When pSILAC is paired with pathway selective perturbations, changes in these kinetic parameters provide a functional readout of pathway dependent stabilization.

A second analytical challenge arises when responses to different degradation pathway perturbations are compared directly. Proteasome and lysosomal inhibition are mechanistically asymmetric interventions and therefore do not necessarily generate comparable effect size distributions. MZB is a covalent and irreversible inhibitor of proteasome catalytic subunits that directly suppresses proteolysis by the UPS^9^. In contrast, BafA1 inhibits the vacuolar H+ ATPase (V-ATPase), disrupts lysosomal acidification, prevents fusion of autophagosome and lysosome at high concentrations or long treatment times, and consequently impairs lysosomal proteolysis and autophagic flux^10, 11^. Thus, the stabilization measured after each treatment reflects not only the biological contribution of the targeted pathway, but also the mechanism, kinetics, and completeness of the pharmacological perturbation.

This pharmacological asymmetry is compounded by differences in pathway usage under basal conditions. The UPS contributes broadly to turnover of soluble intracellular proteins, whereas lysosomal degradation acts on a more selective set of proteins, macromolecular assemblies, and organelles delivered through autophagic and endolysosomal routes. Consequently, even a protein that depends substantially on lysosomal degradation may show a smaller absolute half-life shift after BafA1 treatment than proteins affected by MZB. Direct comparison of absolute half-life changes can therefore conflate pathway preference with perturbation magnitude and systematically favor the perturbation with the broader response distribution. A standardized framework for addressing this dynamic range mismatch in comparative turnover analysis has not, to our knowledge, been established.

We therefore developed a normalization strategy based on within-condition percentile ranks. This transformation places perturbations with different effect size distributions on a common relative response scale while preserving the ordering of protein responses within each condition. The resulting scores report relative pathway preference rather than the absolute contribution of either pathway to protein turnover. We applied this framework to pSILAC measurements following proteasome or lysosome inhibition in HeLa cells and used the resulting degradation preference map to examine the organization of proteome turnover. Of particular interest to us is the partitioning of elements of the proteostasis network between the ALP and the UPS, as we have a long-standing interest in proteostasis, and understanding the dynamics of proteostasis requires an understanding of how the concentrations of proteostasis network (PN) components are controlled. Integration with a curated PN annotation provided a structured biological framework for evaluating this partitioning and revealed coordinated regulation of proteostasis machinery, including lysosomal turnover of proteasome components. Together, this study establishes a general analytical strategy for comparative turnover proteomics and demonstrates its utility for resolving pathway specific proteostasis regulation.

## Results and Discussion

### pSILAC measurement and proteome-wide determination of protein half-lives

To quantify the sensitivity of protein turnover to perturbations of the two major intracellular proteolytic systems, we performed pSILAC experiments in HeLa cells under three conditions: vehicle control (DMSO), proteasome inhibition with MZB, and lysosomal inhibition with BafA1. Cells were cultured in light amino acids before being switched to heavy isotope–labeled medium at the time of treatment. We then followed decay of the preexisting light-labeled protein pool across six time points, allowing degradation kinetics to be estimated for thousands of proteins (Fig. 1A). Representative precursor spectra illustrate the separation of light and heavy isotope envelopes used for kinetic fitting (Fig. 1B).

**Figure 1.**
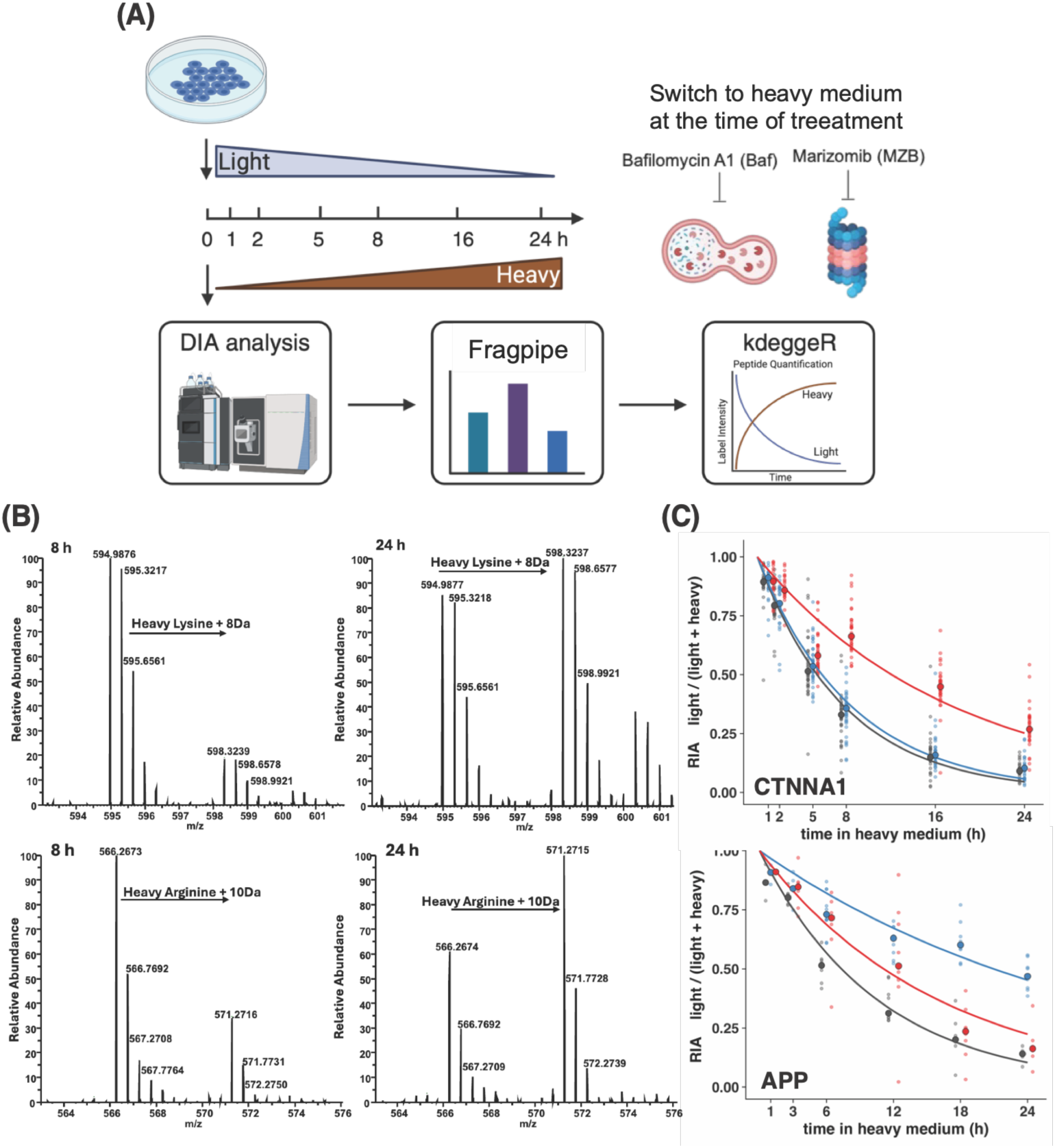
Pulse SILAC design for proteome turnover measurement resolved by degradation. (A) Experimental workflow. HeLa cells cultured in light medium were switched to heavy (^13^C_6_,^15^N_2_-Lys/^13^C_6_,^15^N_4_-Arg) medium at the time of treatment with vehicle (DMSO), marizomib (MZB), or bafilomycin A1 (BafA1). Cells were harvested at six time points during labeling. Decay of the preexisting light pool and accumulation of the newly synthesized heavy pool were quantified by LC-MS/MS. Precursor intensities were extracted with FragPipe and DIA-NN, and degradation rate constants and half-lives were fitted for each protein. (B) Representative precursor isotope envelopes illustrate the range of turnover. Light and heavy forms of the same peptide are separated by +8 Da for heavy lysine or +10 Da for heavy arginine. Spectra are shown as relative abundance versus m/z. **(C)** Representative protein-level k_loss_ decay curves fit from precursor-level relative isotope abundance (RIA) data. Small dots represent RIA values for individual peptide precursors at each time point; large dots represent the corresponding protein-level RIA, and the accompanying line shows the protein-level exponential decay fit used to estimate k_loss_ (black, DMSO; red, MZB; blue, BafA1).

The study consisted of two independent biological experiments, each containing two technical replicates and performed approximately six months apart. To broaden temporal coverage and reduce dependence on a single sampling schedule, the two experiments used offset time series. The first experiment was sampled at 1, 2, 5, 8, 16, and 24 h after the medium switch, whereas the second was sampled at 1, 3, 6, 12, 18, and 24 h.

For each precursor, KdeggeR estimates the loss rate of the pre-existing light-labeled protein pool by fitting the relative isotope abundance of the light peptide (RIA) across the labeling time course (Fig. 1C). Shown on the top panel is CTNNA1, a representative protein preferentially stabilized by proteasome inhibition, showing a slower decline in RIA under MZB (red) relative to BafA1 and DMSO. Shown on the bottom panel is APP, a representative protein preferentially stabilized by lysosomal inhibition, showing a slower decline in RIA under BafA1 (blue) relative to MZB and DMSO. These precursor-level fits are aggregated to protein-level loss rates (k_loss_). The framework then derives degradation rate constants (k_deg_) by correcting k_loss_ for dilution associated with cell division. Because experimentally measured division rates can vary across experiments and treatment conditions, we used the distribution-based correction implemented in KdeggeR, which infers this contribution from the global distribution of protein loss rates. Protein half-lives were calculated from the corrected degradation rate constants as t_1/2_ = ln_(2)_/k_deg_.

Each biological experiment was processed independently through the complete kinetic fitting workflow, and half-lives were calculated before the datasets were integrated. Across both experiments, 5,518 proteins were quantified in at least one condition. Of these, 3,814 proteins yielded high confidence half-life measurements across DMSO, MZB, and BafA1 in both biological experiments. Half-lives from the independently analyzed offset sampling schedules were then averaged to generate the dataset used for comparison across pathways.

### Unequal perturbation responses bias direct comparison across degradation pathways

We first asked whether the turnover responses produced by MZB and BafA1 were sufficiently similar in magnitude to support direct comparison. They were not. The two inhibitors produced markedly different proteome wide half-life distributions (Fig. 2A). Median protein half-life increased from 23.1 h under control conditions to 34.4 h after MZB treatment, a median increase of 11.3 h. In contrast, BafA1 increased the median half-life from 23.1 to 24.6 h, a shift of only 1.5 h. The median stabilization produced by MZB was therefore approximately 7.5-fold greater than that produced by BafA1.

**Figure 2.**
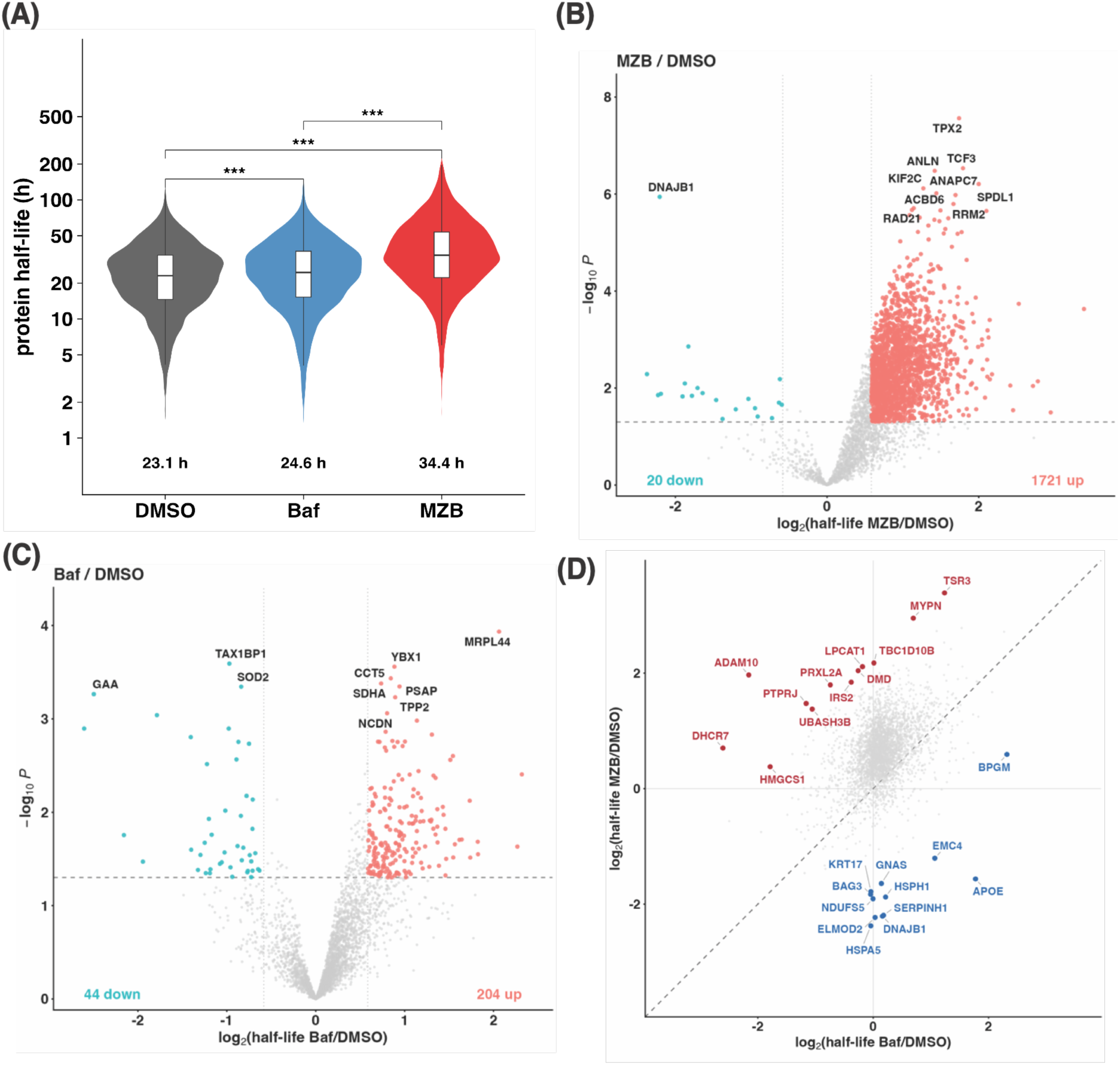
Proteasome and lysosome inhibition produce turnover responses with markedly different dynamic ranges. (A) Distribution of protein half-lives under each condition for 3,814 proteins with high confidence fits in all three conditions. Violins show the full distribution on a log scale, and boxes indicate the interquartile range and median. Protein half-lives were compared between conditions using two-sided Wilcoxon signed-rank tests, paired by protein, p < 0.001 (***). (B, C) Volcano plots of half-life change for MZB versus DMSO (B) and BafA1 versus DMSO (C). Each point represents one protein. The log2 half-life fold change is plotted against minus log10 P value. Vertical lines mark the fold change cutoff, and the horizontal dotted line marks p = 0.05. Proteins passing both thresholds are colored red for stabilization or blue for destabilization. The most extreme proteins are labeled. (D) Direct comparison of unnormalized effect sizes under MZB and BafA1. The diagonal marks equal response. The point cloud is displaced toward the MZB axis, illustrating the mismatch in dynamic range that motivates rank normalization.

This difference should not be interpreted as a direct quantitative measure of the relative contributions of the UPS and ALP to protein degradation. The observed effect size reflects both biological pathway usage and the mechanism, kinetics, and completeness of each pharmacological perturbation. MZB directly and irreversibly inhibits proteasome catalytic activity, whereas BafA1 suppresses lysosomal proteolysis indirectly by disrupting V-ATPase dependent acidification. The two treatments therefore generate responses with intrinsically different dynamic ranges. When raw half-life changes are compared directly, the broader MZB distribution dominates the analysis, so proteins can appear preferentially dependent on the proteasome simply because MZB produces a larger absolute stabilization response.

Two representative proteins illustrate this problem. GAPDH, which can undergo lysosomal degradation through chaperone-mediated autophagy^13^, had a basal half-life of 25.8 h that increased to 29.8 h with BafA1 (Δt_1/2_ = 4.0 h) but to 70.4 h with MZB (Δt_1/2_ = 44.6 h). EGFR, which is cleared through the endolysosomal system^14^, had a basal half-life of 9.3 h that increased to 13.0 h with BafA1 (Δt_1/2_ = 3.7 h) and to 26.5 h with MZB (Δt_1/2_ = 17.2 h). If pathway assignment were based only on the magnitude of the half-life shift, both proteins would appear strongly proteasome preferential despite established lysosomal routes contributing to their turnover. These examples illustrate why absolute effect size alone is insufficient for comparison across pathways.

Direct comparison of the raw fold changes further revealed a pronounced asymmetry, with the proteome wide point cloud displaced toward the MZB axis (Fig. 2B-D). These observations show that absolute stabilization cannot be interpreted on a common numerical scale across the two perturbations. A larger half-life change under MZB than under BafA1 does not necessarily establish preferential UPS degradation because the expected response magnitude differs systematically between the interventions. Comparison across pathways therefore requires each protein to be evaluated relative to the response distribution produced by the corresponding perturbation.

Gene ontology enrichment analysis further demonstrated that the two perturbations produced distinct functional responses. Proteins stabilized by BafA1 were enriched for mitochondrial envelope and inner membrane terms, consistent with lysosome-dependent turnover of mitochondrial constituents. In comparison, proteins stabilized by MZB were enriched for ribonucleoprotein complex and cell adhesion terms, whereas proteins with reduced apparent half-lives were enriched for protein folding and unfolded protein response pathways (Fig. S1). The latter observation requires caution because proteasome inhibition induces proteotoxic stress and can alter synthesis of chaperones and other stress responsive proteins. Apparent changes in turnover may therefore reflect adaptive proteome remodeling in addition to altered degradation. These enrichment patterns reinforce the conclusion that MZB and BafA1 elicit biologically distinct proteome-wide responses and further motivate a relative, rather than absolute, framework for comparison.

### Within-condition rank normalization enables comparison across pathways with unequal response ranges

We therefore sought a transformation that would preserve the relative ordering of protein responses within each perturbation without requiring the absolute effect size distributions of MZB and BafA1 to be equivalent. For each treatment, we calculated the log2 fold change in half-life relative to DMSO and ranked proteins according to the magnitude of this response. Ranks were converted to percentiles using the Hazen plotting position, p = (r - 0.5)/n, where r is the rank and n is the number of proteins. Tied values were assigned the average rank. The resulting scores are centered around 0.5 and lie within the open interval (0, 1), where values below 0.5 indicate stronger stabilization by MZB treatment; values above 0.5 indicate stronger stabilization by BafA1 treatment. For the averaged dataset (n = 3,814), percentile values ranged from 0.000131 to 0.999869.

This transformation changes the question being asked. Direct comparison asks whether a protein is stabilized by a larger absolute amount under MZB or BafA1. Rank normalization instead asks how strongly that protein responds to each perturbation relative to all other proteins measured under the same condition. For example, a protein in the 90th percentile after BafA1 treatment is among the most strongly lysosome-sensitive proteins in the measured proteome even if its absolute increase in half-life is smaller than that of a protein occupying a lower percentile after MZB treatment.

Percentile transformation therefore places the two perturbations on a common relative response scale while preserving their internal response order. Plotting the normalized MZB response against the normalized BafA1 response places each protein in a two-dimensional degradation preference space (Fig. 3A). The diagonal (dashed line) represents equal relative sensitivity to the two perturbations, not equal absolute stabilization. Displacement from the diagonal indicates a stronger response relative to the distribution generated by one inhibitor or the other. Accordingly, rank normalization reports relative degradation pathway preference rather than the absolute contribution of either pathway to protein turnover. This distinction is central to interpretation of the normalized map.

**Figure 3.**
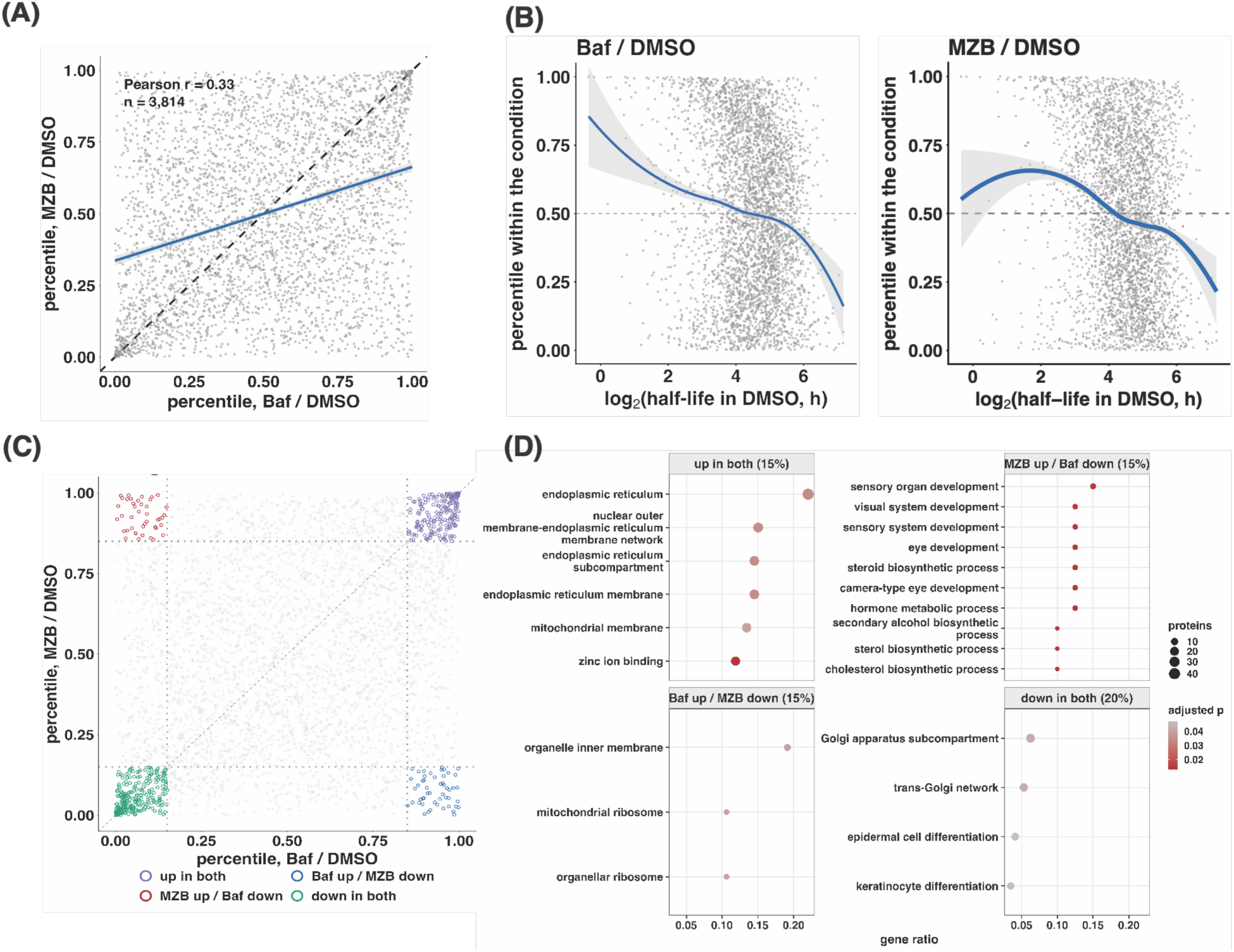
Percentile rank normalization places both perturbations on a common scale and resolves four degradation preference regimes. (A) Percentile response under MZB versus BafA1 for all 3,814 proteins. Each axis represents the percentile rank of the log2 half-life fold change within that treatment, calculated as p = (rank - 0.5)/n using the Hazen plotting position. Both marginal distributions are therefore uniform on (0, 1) by construction. The blue line is the ordinary least squares fit, and the dashed line is the identity diagonal. (B) Percentile response within each condition plotted against log2 basal half-life in DMSO. Blue lines are local regression fits with 95% confidence bands, and the dotted line marks the median percentile of 0.5. (C) Degradation preference map with corner regions highlighted. Dotted lines indicate the percentile thresholds used to define four regimes: high response to both perturbations, high MZB and low BafA1 response, high BafA1 and low MZB response, and low response to both perturbations. (D) Gene ontology overrepresentation within each corner. Dot size represents the number of proteins assigned to the term, color indicates the Benjamini-Hochberg adjusted p value, and the x axis shows gene ratio.

The normalized MZB and BafA1 responses were positively associated (Spearman rank correlation ρ = 0.33), indicating that there is a general tendency for proteins to respond similarly to both perturbations; nevertheless, many show clear relative preference. Because percentile scores are monotonic transformations of the original response ranks within each condition, Spearman correlation is the appropriate measure of association between the two normalized axes.

We next related percentile response to basal half-life (Fig. 3B). Under BafA1, the relationship was approximately monotonic: proteins with shorter basal half-lives tended to occupy higher stabilization percentiles, and the percentile response declined as basal half-life increased. The pattern under MZB was nonmonotonic. The highest percentiles were concentrated among proteins with intermediate basal stability, whereas long-lived proteins were distributed across both higher and lower portions of the ranking. These distinct response architectures were less apparent in the raw fold change analysis and further support use of a relative scale for comparison across pathways.

### The normalized framework resolves four degradation response regimes

We next used the normalized map to identify broad patterns of pathway preference. Four response regimes were defined by the corners of the two-dimensional preference space (Fig. 3C). Shown in Fig. 3D, proteins strongly stabilized by both inhibitors (or “up”) were enriched for endoplasmic reticulum, nuclear outer membrane and ER membrane network, ER membrane, and mitochondrial membrane compartments, together with zinc ion binding functions. Proteins with low stabilization ranks under both perturbations (or “down”) were enriched for Golgi subcompartments and trans Golgi network terms, as well as epidermal cell and keratinocyte differentiation. Proteins preferentially stabilized by proteasome inhibition were enriched for sensory and eye development terms and for sterol and cholesterol biosynthesis. In contrast, proteins preferentially stabilized by lysosomal inhibition were strongly enriched for organelle inner membrane, mitochondrial ribosome, and organellar ribosome terms, consistent with lysosome-dependent turnover of membrane and mitochondrial components^15, 16^. These patterns reveal functional organization that is difficult to resolve from the unnormalized comparison and demonstrate the utility of the rank framework for identifying relative pathway structure despite unequal perturbation magnitudes.

Rank normalization removes differences in the global response distributions, but it cannot eliminate protein-specific effects caused by altered synthesis or compensatory signaling. This limitation is particularly relevant for proteasome inhibition, which can activate the integrated stress response (ISR) and remodel global translation^17, 18^. We therefore examined the preference ranks of ISR-associated proteins. These proteins did not systematically cluster within a single region of the degradation preference map (Fig. S2), arguing against the ISR as the dominant driver of pathway assignment. Nevertheless, orthogonal perturbations will be important for distinguishing direct degradation effects from secondary stress responses.

### Proteostasis network annotation reveals coherent pathway preferences

Having established the normalized preference map, we next asked whether its assignments were biologically coherent at the level of organized proteostasis systems. We integrated the dataset with a curated proteostasis network (PN) annotation framework^19^, which classifies proteins into nine major branches: mitochondrial proteostasis, ER proteostasis, cytonuclear proteostasis, nuclear proteostasis, cytosolic translation, proteostasis regulation, extracellular proteostasis, the autophagy-lysosome pathway, and the ubiquitin-proteasome system. Of the 3,814 proteins with high confidence half-life measurements in all three conditions, 1,648 carried at least one PN branch annotation. The annotated set included 539 UPS, 427 ALP, 384 translation, 145 ER proteostasis, 107 cytonuclear, 81 nuclear, 49 mitochondrial, 9 PN regulation, and 5 extracellular annotations (Fig. 4A). Branch counts are not mutually exclusive because individual proteins can carry more than one annotation.

**Figure 4.**
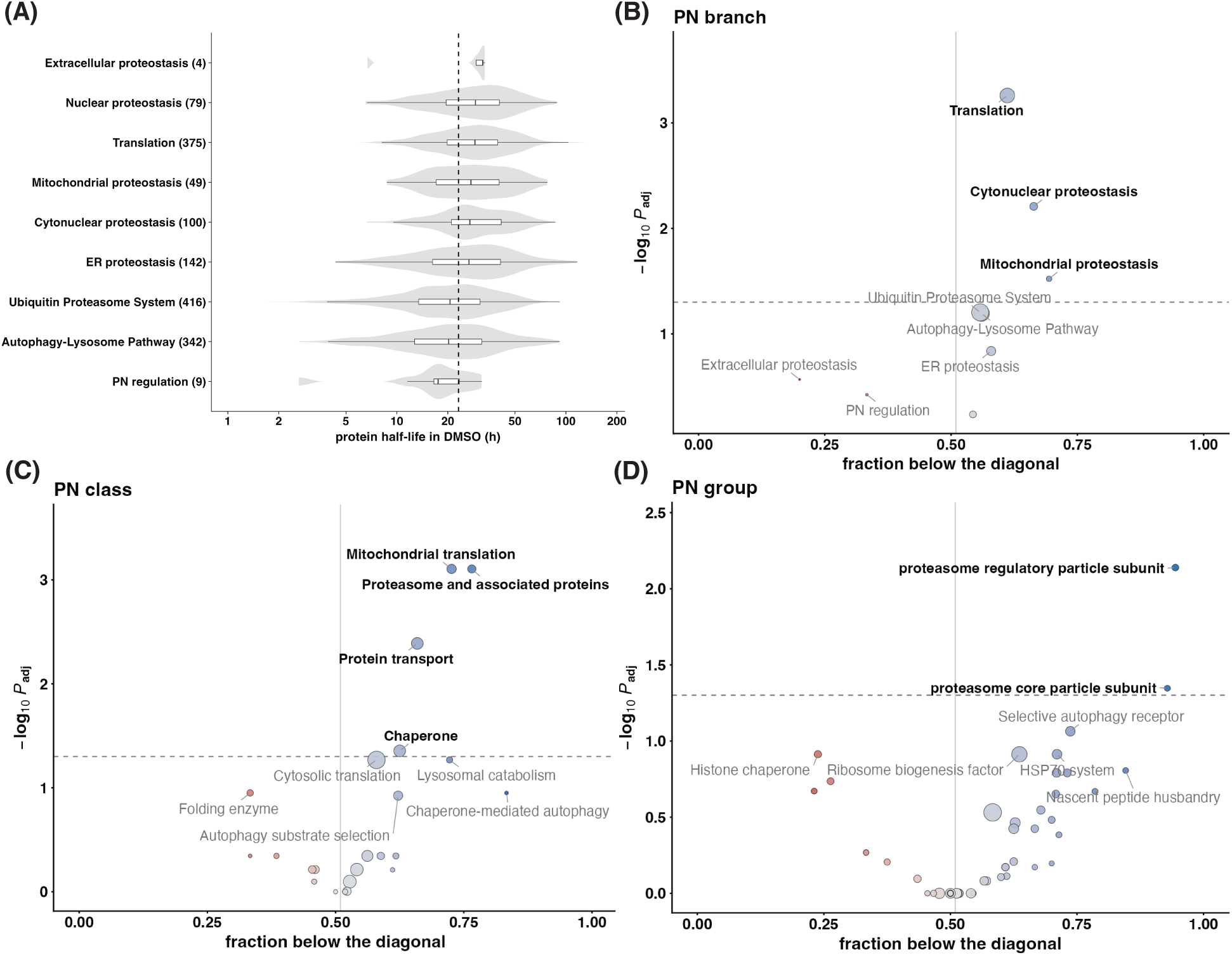
Proteostasis network annotation reveals degradation preferences at the branch, class, and group levels. (A) Basal half-life distribution in DMSO by proteostasis network branch^19^. Violins show the full distribution on a log scale with overlaid boxplots. The number of quantified proteins in each branch is given in parentheses, and the vertical dashed line marks the proteome-wide median. (B to D) Sign tests for degradation pathway preference at the branch (B), class (C), and group (D) levels. Bubble color indicates the direction of the group level sign test, with blue indicating ALP preference and red indicating UPS preference. The vertical line marks a fraction of 0.5, and the horizontal dotted line marks p = 0.05. Sets reaching statistical significance are labeled with black text.

Basal half-life distributions differed across PN branches (Fig. 4A). Nuclear, mitochondrial, cytonuclear, and ER proteostasis proteins, together with proteins involved in translation, generally had median half-lives above the proteome-wide median. In contrast, proteins assigned to the ALP and UPS tended to turn over more rapidly^20^. The extracellular and PN regulation branches contained too few quantified proteins for firm interpretation. Overall, these patterns suggest that components of the degradation machinery can be remodeled more rapidly than many of the structural and organellar clients they maintain.

We then tested whether individual PN branches showed systematic pathway preference. Across all PN-annotated proteins, there was a modest but significant overall bias toward the ALP-preferential side of the diagonal (759 of 1,326 proteins, ∼57%). For each branch, class, and group, we applied the sign test to the number of member proteins above versus below the diagonal, with proteins above classified as UPS preferential (red) and below as ALP preferential (blue) (Fig. 4 B-D). At the branch level, translation, cytonuclear proteostasis, and mitochondrial proteostasis showed a significant ALP preference; the ubiquitin–proteasome system and autophagy–lysosome pathway branches showed the same directional trend but with higher p_adj_ values that did not reach significance(Fig. 4B). The remaining branches were distributed closer to the diagonal, indicating more balanced relative sensitivity to the two degradation systems (Figure S3).

Because each broad PN branch contains mechanistically diverse protein classes, we repeated the sign test at the finer class and group levels. At the class level, four classes reached significance, all showing preferential sensitivity to lysosomal inhibition: mitochondrial translation, proteasome and associated proteins, protein transport, and chaperones(Fig. 4C). Cytosolic translation, lysosomal catabolism, and chaperone-mediated autophagy showed the same directional trend with higher, non-significant p_adj_ values; these are labeled in Fig. 4C to indicate the trend rather than a confirmed effect. Folding enzymes showed a bias toward proteasome-preferential sensitivity, but with a p_adj_ value that did not reach significance. At the group level, two groups showed significant p_adj_ values, both corresponding to proteasome subcomplexes and both preferentially sensitive to lysosomal inhibition: proteasome regulatory particle subunits and proteasome core particle subunits (Fig. 4D). Several additional groups, including selective autophagy receptors, the HSP70 system, ribosome biogenesis factors, nascent peptide husbandry factors, and histone chaperones, showed directional trends consistent with lysosomal or proteasomal preference, respectively, with higher p_adj_ values that did not reach significance. Individual proteins within each class and group are shown in the corresponding scatter plots (Figs. S4 and S5). Broadly speaking, our results suggest that many proteostasis pathways are dependent on the ALP for their turnover. This observation is consistent with many of these pathways relying on complexes that are degraded as a unit (e.g., the proteasome and the ribosome), or existing in organelles that are degraded through selective autophagy.

Several of these assignments provide an internal biological test of the normalized framework. Most notably, both proteasome core particle and regulatory particle subunits emerged among the groups with the strongest relative sensitivity to lysosomal inhibition (Fig. 4D), consistent with proteaphagy, the selective autophagic turnover of proteasomes^21–23^, being an important pathway for the recycling of proteasomes. Coordinated behavior across multiple subunits of the same macromolecular complex is particularly informative because it indicates that the analysis captures pathway-level organization rather than isolated canonical examples. The relative lysosomal sensitivity of chaperone-mediated autophagy components and lysosomal catabolic proteins is also consistent with their residence in, or delivery to, the lysosomal system. Together, these results show that PN annotation provides a biologically structured framework for interpreting degradation preference maps and reveals coordinated regulation of proteostasis machinery that is less apparent from conventional gene ontology analysis alone.

### mTOR inhibition promotes autophagic degradation of proteasomes

The PN analysis identified proteasome components as one of the groups with the strongest relative sensitivity to lysosomal inhibition, providing a direct biological prediction from the normalized turnover map. Proteaphagy, the selective autophagic degradation of proteasomes, has been observed in several systems but remains less well characterized in mammalian cells than other forms of selective autophagy. We therefore tested whether activation of autophagy would reduce proteasome abundance. Treatment with the mTOR inhibitor Torin1^24^ (250 nM, 24 h) decreased the abundance of the proteasome regulatory subunits PSMC2 and PSMD2 by immunoblotting (Fig. 5A), consistent with enhanced proteasome degradation during autophagy activation.

**Figure 5.**
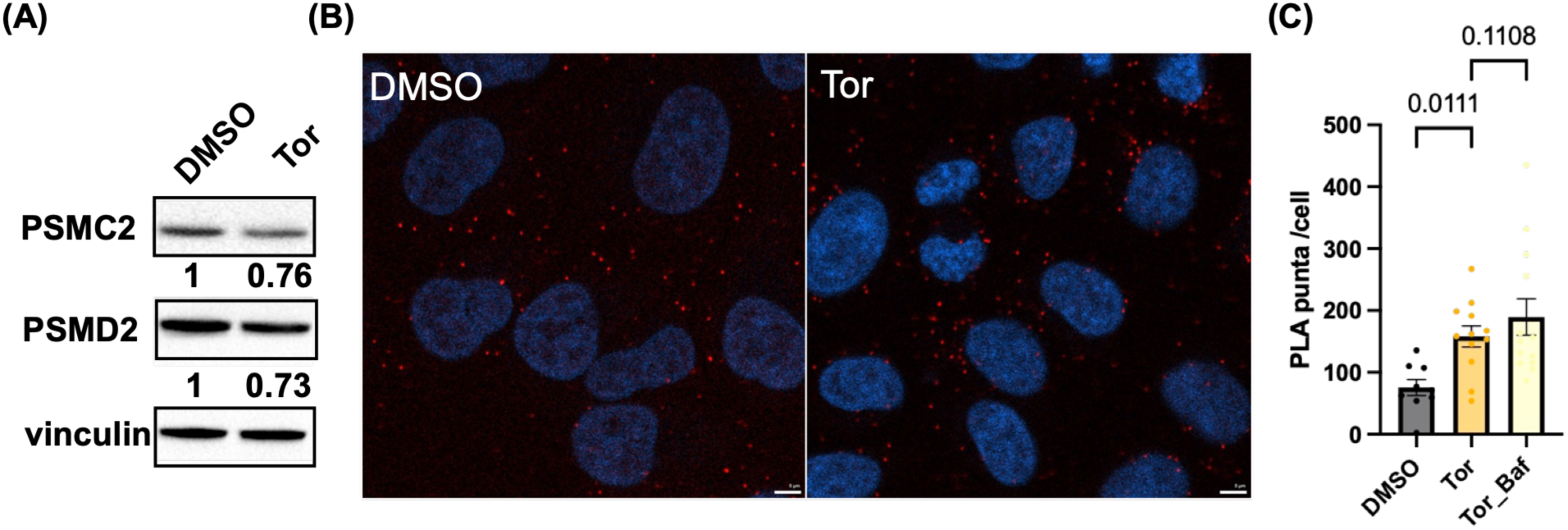
mTOR inhibition promotes proteasome degradation and increases proximity between proteasomes and LC3. (A) Immunoblot of the proteasome regulatory subunits PSMC2 and PSMD2 in HeLa cells treated with DMSO or Torin1 (Tor; 250 nM, 24 h). (B) Representative confocal images of proximity ligation assay (PLA) signal between the proteasome subunit PSMB5 and the autophagy marker LC3 in HeLa cells treated with DMSO or Torin1. Nuclei are counterstained with DAPI. Scale bar, 5 µm. (C) Quantification of PLA puncta per cell across DMSO, Torin1, and Torin1 plus bafilomycin A1 treatment conditions. Each dot represents one image, and bars represent mean ± SEM. Statistical significance was assessed using an unpaired two tailed t test.

To determine whether proteasomes were recruited to autophagic structures, we performed proximity ligation assays (PLA) between the proteasome subunit PSMB5 and the autophagy marker LC3 (Fig. 5B). Few LC3 and PSMB5 PLA puncta were detected under basal conditions. Torin1 treatment markedly increased the number of puncta, indicating increased proximity between proteasomes and LC3-positive structures during autophagy activation. Combining BafA1 with Torin1 produced a further increase in PLA puncta (Fig. 5C), consistent with accumulation of proteasome-containing autophagic intermediates when lysosomal degradation is impaired. These orthogonal observations support the biological interpretation of the normalized pSILAC map and provide evidence that mammalian proteasomes undergo autophagic turnover when autophagic flux is increased.

## Conclusion

This study establishes a rank normalization framework for comparing protein turnover responses across perturbations that generate substantially different effect size distributions. Applied to pSILAC measurements of proteasome and lysosome inhibition in HeLa cells, the framework converts absolute stabilization into relative responses within each condition. Integration of these responses with annotation systems like Gene Ontology and the proteostasis network annotation then enables systematic assessment of relative degradation pathway preference across the proteome. Our analysis suggests that proteostasis network components, especially those that exist as parts of complexes or in organelles, are generally more ALP sensitive that the rest of the proteome. Importantly, the normalized scores should be interpreted as measures of relative sensitivity to each perturbation rather than as estimates of the absolute fraction of protein turnover mediated by the UPS or ALP.

Several limitations of the underlying turnover measurement should be considered. Residue-specific metabolic labeling reports protein loss imperfectly because light lysine and arginine released during degradation of the preexisting proteome can reenter the intracellular precursor pool and be reused for synthesis. As a result, the light signal can decay more slowly than the corresponding protein pool, leading to systematic overestimation of fitted half-lives^25, 26^. Labeling with the methionine analog azidohomoalanine has been proposed as an alternative because the analog is not efficiently recycled, but this strategy also has limitations, including incomplete incorporation and measurable effects on the transcriptome and proteome^27, 28^.

The unequal response distributions observed here likely arise from a combination of biological, pharmacological, and adaptive factors. The UPS and ALP differ intrinsically in their contributions to basal protein turnover, while MZB and BafA1 perturb these systems through distinct molecular mechanisms with different kinetics and degrees of inhibition. In addition, each treatment can trigger secondary responses. Proteasome inhibition activates the integrated stress response and can remodel translation, whereas lysosomal inhibition can disrupt nutrient recycling and lysosome-dependent signaling. Rank normalization addresses the analytical consequence of unequal global response distributions, but it does not distinguish among these underlying causes or eliminate protein-specific secondary effects. Orthogonal perturbations, including genetic depletion such as ATG5 or ATG7 knockout and inhibitors acting at different steps of each pathway, will therefore be important for validating individual pathway assignments.

Although we demonstrate the approach using proteasome and lysosome inhibition in HeLa cells, the analytical principle should apply broadly to turnover datasets in which experimental perturbations generate unequal response distributions. The framework can be extended to genetic perturbations, alternative pathway selective inhibitors, and targeted protein degradation strategies, including PROTAC^29^ and AUTOTAC^30^, provided that the resulting scores are interpreted as relative responses within each perturbation. More broadly, combining degradation-resolved proteomics with structured proteostasis annotations offers a quantitative strategy for examining how proteome turnover is partitioned across degradation pathways and for identifying coordinated regulation of proteostasis machinery in cellular and disease relevant systems.

## Materials and Methods

### Cell culture and drug treatments

HeLa cells were maintained in Dulbecco’s Modified Eagle Medium (DMEM) supplemented with 10% fetal bovine serum (FBS), 100 U/mL penicillin, and 100 µg/mL streptomycin at 37°C in a humidified atmosphere containing 5% CO2. For proteasome inhibition, cells were treated with marizomib (MZB); for lysosomal inhibition, cells were treated with bafilomycin A1 (BafA1). Vehicle control cells received an equivalent volume of DMSO. All treatments were performed in biological triplicate.

### Pulse SILAC labeling

HeLa cells were cultured in DMEM containing light (unlabeled) L-arginine and L-lysine supplemented with 10% dialyzed FBS for at least six cell doublings to ensure complete proteome labeling. At the start of each experiment, the medium was switched to heavy SILAC medium containing ^13^C_6_,^15^N_4_-L-arginine (Arg10) and ^13^C_6_,^15^N_2_-L-lysine (Lys8) supplemented with 10% dialyzed FBS, and cells were simultaneously treated with DMSO, marizomib, or bafilomycin A1. Cell pellets were collected at multiple time points following the medium switch to monitor the decay of pre-existing light-labeled proteins. Three biological replicates were performed for each condition.

### Sample preparation for mass spectrometry

Cells were lysed in a buffer containing 8 M urea, 75 mM NaCl, 50 mM HEPES (pH 7.5), cOmplete™ Mini EDTA-free protease inhibitor cocktail (Sigma-Aldrich, #11836170001), and PhosSTOP phosphatase inhibitor cocktail (Sigma-Aldrich, #04906837001). Lysates were incubated on ice for at least 15 minutes with occasional vortexing to ensure complete lysis, then cleared by centrifugation at 13,000 rpm for 15 minutes at 4°C. Protein concentration in the resulting supernatant was determined by BCA assay (Pierce™ BCA Protein Assay Kit, Thermo Fisher Scientific, #23227). Equal amounts of protein from each sample were reduced with 5 mM Tris(2-carboxyethyl)phosphine (TCEP) for 30 minutes at room temperature, alkylated with 14 mM iodoacetamide (IAA) for 30 minutes at room temperature in the dark.

Proteins were processed using a single-pot, solid-phase-enhanced sample preparation (SP3) workflow with Sera-Mag carboxylate-modified magnetic beads (Cytiva). A 1:1 mixture of hydrophilic and hydrophobic Sera-Mag beads was added to each sample at a bead-to-protein ratio of approximately 5–10:1 (µg:µg), followed immediately by an equal volume of 100% ethanol to induce protein binding. Samples were incubated at 24°C with shaking (1,000 rpm) for 10 minutes, and beads were then collected on a magnetic rack and washed three times with 80% ethanol. Bead-bound protein was resuspended in 200 mM EPPS (pH 8.0) and digested with Trypsin/Lys-C Mix (Thermo Fisher Scientific, #A40009) at a 1:100 enzyme-to-protein ratio overnight at 37°C with shaking. Following digestion, an additional aliquot of 200 mM EPPS was added to the beads and incubated for 10 minutes to elute residual peptides, and the combined eluates were collected on a magnetic rack to remove residual beads.

Peptide digests were acidified to a final concentration of 0.1–2% trifluoroacetic acid (TFA) and desalted using Pierce™ Peptide Desalting Spin Columns (Thermo Fisher Scientific) according to the manufacturer’s protocol. Desalted peptides were dried by vacuum centrifugation, resuspended in 0.1% formic acid, and peptide concentration was determined by absorbance at 205 nm. Approximately 0.5–1 µg of peptide was injected for each LC–MS/MS analysis.

### Liquid chromatography–tandem mass spectrometry (LC–MS/MS)

Peptides were separated by nanoflow liquid chromatography using a Vanquish Neo UHPLC system (Thermo Fisher Scientific) coupled online to a FAIMS Pro interface and an Orbitrap Exploris 480 mass spectrometer (Thermo Fisher Scientific) operated in data-independent acquisition (DIA) mode. Samples were loaded and separated on a 50 cm EASY-Spray™ HPLC column (75 µm inner diameter, ES903, Thermo Fisher Scientific) using a trap-and-elute configuration at a flow rate of 0.35 µL/min over a 120-minute gradient. The mobile phase consisted of water (solvent A) and 80% acetonitrile (solvent B), and the gradient was ramped from 3% to 22% B over 78 minutes, then to 37% B by 112 minutes, followed by a column wash to 99% B before re-equilibration. Ions were generated by nanoelectrospray ionization (positive mode, 1,900 V) with an ion transfer tube temperature of 300°C. FAIMS separation was performed in standard resolution mode with a total carrier gas flow of 4 L/min, and data were acquired using two alternating compensation voltages (CV, –40 V and –60 V), each paired with a full MS scan followed by a corresponding DIA scan (four total experiments per cycle). Full MS scans were acquired at each CV over a scan range of 380–980 m/z at an Orbitrap resolution of 120,000, with a normalized AGC target of 300% (absolute target 3.0 × 10⁶), automatic maximum injection time, and profile-mode data acquisition. DIA scans were acquired across the same precursor mass range (380–980 m/z) using sequential 14 m/z isolation windows with 0.5 m/z overlap (43 scan events per CV), fragmented by higher-energy collisional dissociation (HCD) at a normalized collision energy of 28%. DIA fragment spectra were acquired at an Orbitrap resolution of 15,000 over a scan range of 145–1,450 m/z, with a normalized AGC target of 1000% (absolute target 1.0 × 10⁶), automatic maximum injection time, and centroid-mode data acquisition.

### Proteomics data analysis and half-life determination

Raw DIA mass spectrometry data were processed using FragPipe (version 22.0). Spectral libraries were generated directly from the DIA data using MSFragger together with EasyPQP, and retention time and fragmentation predictions were refined using MSBooster with DIA-NN-based deep learning models. The resulting project-specific spectral libraries were used for quantification with DIA-NN, run in multiplexed (plexDIA) mode to resolve light and heavy SILAC peptide channels. Light and heavy labels were defined as unmodified lysine and arginine (K0/R0) and heavy lysine (+8.014199 Da, Lys8) and arginine (+10.008269 Da, Arg10), respectively, consistent with the pSILAC labeling scheme. Peptide and protein identifications were filtered at a 1% false discovery rate at both the MSFragger/Percolator/ProteinProphet level and the DIA-NN precursor q-value level, and quantification was performed using DIA-NN’s QuantUMS (high-accuracy) algorithm. Searches were performed against the reviewed UniProt human reference proteome (UP000005640) appended with common contaminants and reversed decoy sequences. Trypsin was specified as the digestion enzyme with up to one missed cleavage, carbamidomethylation of cysteine was set as a fixed modification, and oxidation of methionine and protein N-terminal acetylation were set as variable modifications, in addition to the heavy SILAC labels described above.

Protein turnover rates were determined from the resulting light- and heavy-channel peptide quantification using the KdeggeR framework. Briefly, KdeggeR estimates the loss rate of the pre-existing light-labeled protein pool by fitting the decay of relative light isotope abundance across the labeling time course and derives degradation rate constants (kdeg) after correcting for dilution associated with cell division. Rather than supplying an experimentally measured division rate, which can vary across experiments and treatment conditions, we used the distribution-based correction implemented in KdeggeR, in which the correction is inferred from the global distribution of protein loss rates. Protein half-lives were then calculated from the corrected degradation rate constants as t_1/2_ = ln_(2)_/k_deg_.

Each biological replicate was processed independently through the complete search-and- quantification-to-kinetic-fitting workflow, and half-lives were calculated separately before integration. Half-lives from the two independently analyzed, offset-schedule experiments were then averaged to generate the final dataset used for cross-pathway comparison. Proteins were retained for downstream analysis only when quantified with acceptable fit quality in at least two of the independent replicates, ensuring that subsequent pathway assignments were based on robust kinetic measurements rather than poorly constrained fits.

### Proximity ligation assay (PLA)

Proximity ligation assays were performed using the Duolink® In Situ Detection Reagents Far Red kit (Sigma-Aldrich) according to the manufacturer’s protocol. HeLa cells were seeded on glass coverslips and treated with DMSO, Torin1 (250 nM, 24 h), or Torin1 in combination with bafilomycin A1 (BafA1) as indicated. Cells were fixed, permeabilized, and blocked using Duolink blocking solution, then incubated overnight at 4°C with primary antibodies against PSMB5 (Proteintech, 67959, 1:100) and LC3B (CST, 2775, 1:100), raised in different host species. Samples were subsequently incubated with species-specific Duolink PLUS and MINUS PLA probes, followed by ligation and rolling-circle amplification performed according to the Duolink Far Red protocol. Coverslips were mounted using Duolink mounting medium containing DAPI to visualize nuclei.

Confocal images of PLA assay was acquired on Zeiss Cell Discoverer 7 microscopy (50x magnification with numeric aperture of 1.2). The acquired images were analyzed using Arivis (v4.20, Zeiss). PLA puncta and neucleus segmentation were performed with the machine-learning module of the analysis pipeline, using training datasets generated by manually annotating areas of interest at random. The analysis pipeline was applied in batch mode to generate quantification data for each well. The number of PLA puncta and neucleus results calculated and exported as number of PLA puncta per neucleus per well.

### Statistical analysis

Statistical analysis and data visualization were performed using R or Python developed with the assistance of Claude (Anthropic). The sign test was used to evaluate bias in the distribution of proteostasis network branch members relative to the diagonal in the degradation preference space. Enrichment analyses used multiple testing correction by the Benjamini–Hochberg method. All resulting figures were independently verified by the authors against the underlying data for accuracy.

## Data availability

The mass spectrometry proteomics data have been deposited to the ProteomeXchange Consortium via the PRIDE partner repository. Processed protein half-life measurements and normalized degradation preference scores for all quantified proteins are provided as Supplementary Tables.

## Acknowledgments

This work is supported by R01AG073418 and the Freedom Together Foundation. We greatly appreciate Emily P. Bentley’s expert editorial feedback.

---

**Figure S1.**
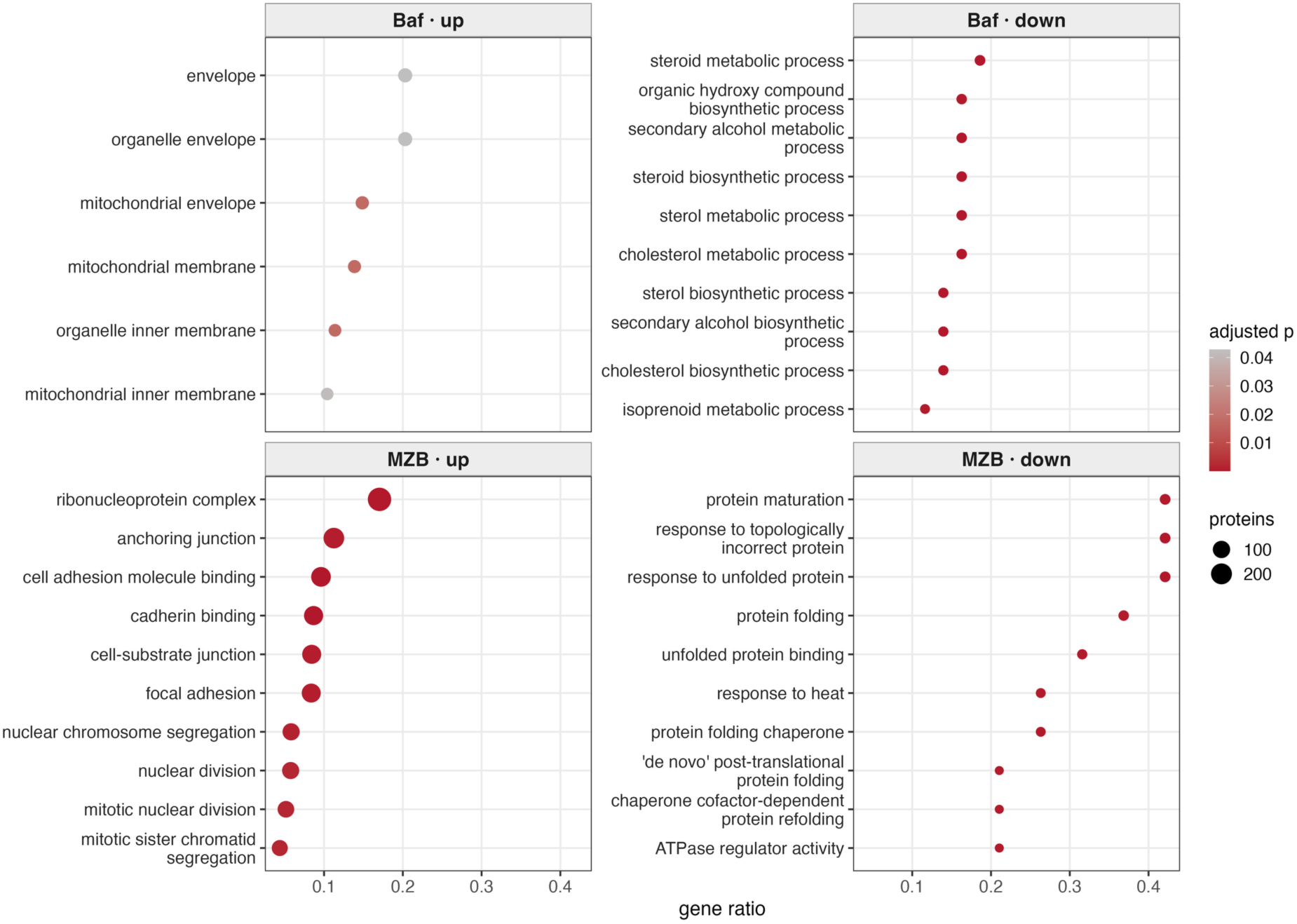
Gene ontology overrepresentation among proteins with altered apparent half-life under each perturbation. Terms enriched among proteins stabilized (up) or destabilized (down) by BafA1 and MZB were tested separately against the quantified proteome. Dot size represents the number of annotated proteins, color indicates the Benjamini-Hochberg adjusted p value, and the x axis shows gene ratio.

**Figure S2.**
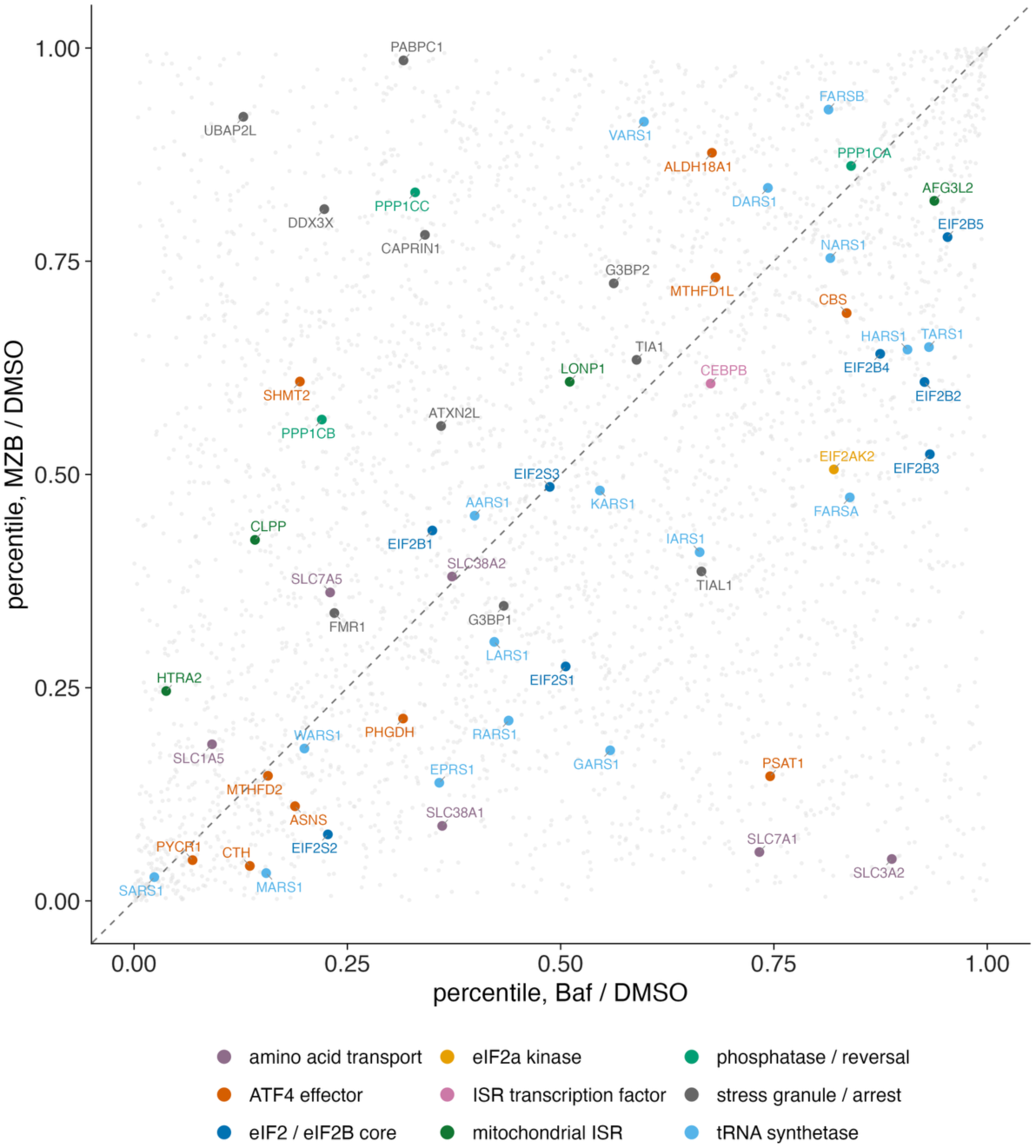
Integrated stress response proteins do not occupy a single region of the degradation preference map. Percentile responses under MZB and BafA1 are shown for manually curated ISR associated proteins overlaid on the full quantified proteome in grey. Points are grouped by functional category and labeled with gene symbols. The dashed line is the identity diagonal.

**Figure S3.**
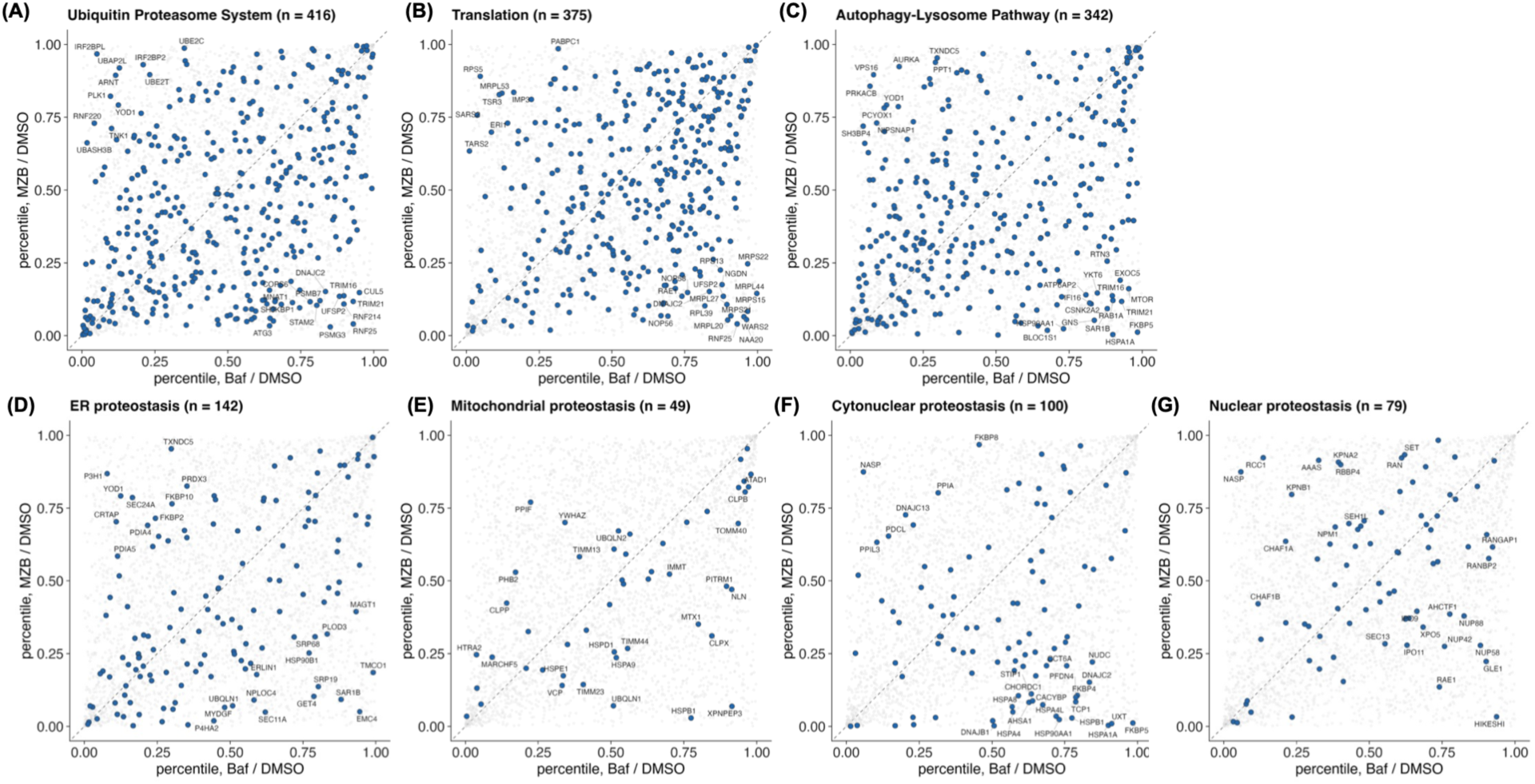
Degradation preference maps for individual proteostasis network branch. Percentile responses under MZB and BafA1 are shown for members of each PN class (A to G) overlaid on the full quantified proteome in grey. The class name and number of quantified members are given in each panel title. Point color indicates the direction of the class level sign test, with blue indicating ALP preference and red indicating UPS preference. The dashed line is the identity diagonal, and individual proteins are labeled with gene symbols.

**Figure S4.**
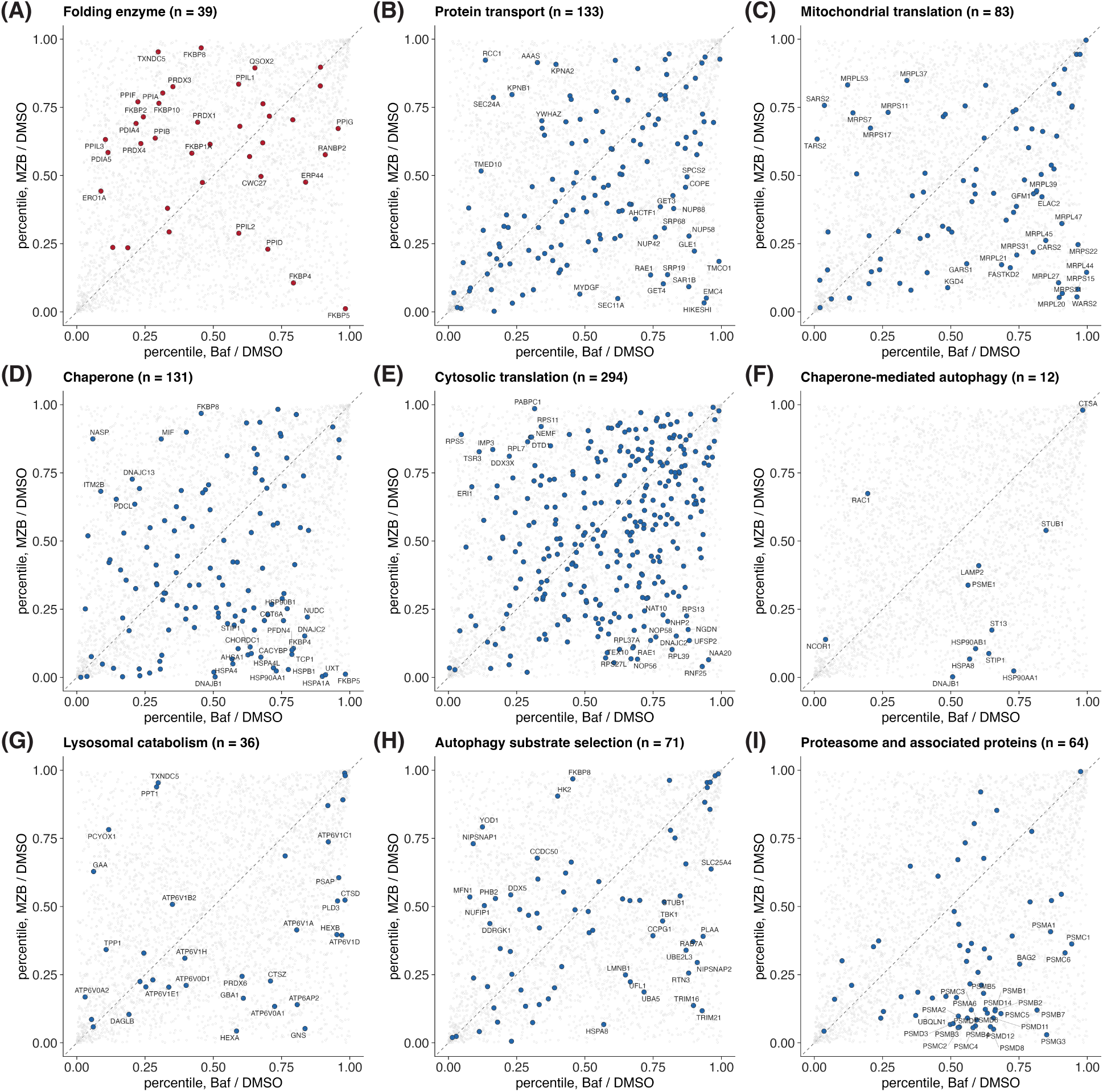
Degradation preference maps for individual proteostasis network classes. Percentile responses under MZB and BafA1 are shown for members of each PN class (A to I) overlaid on the full quantified proteome in grey. The class name and number of quantified members are given in each panel title. Point color indicates the direction of the class level sign test, with blue indicating ALP preference and red indicating UPS preference. The dashed line is the identity diagonal, and individual proteins are labeled with gene symbols.

**Figure S5.**
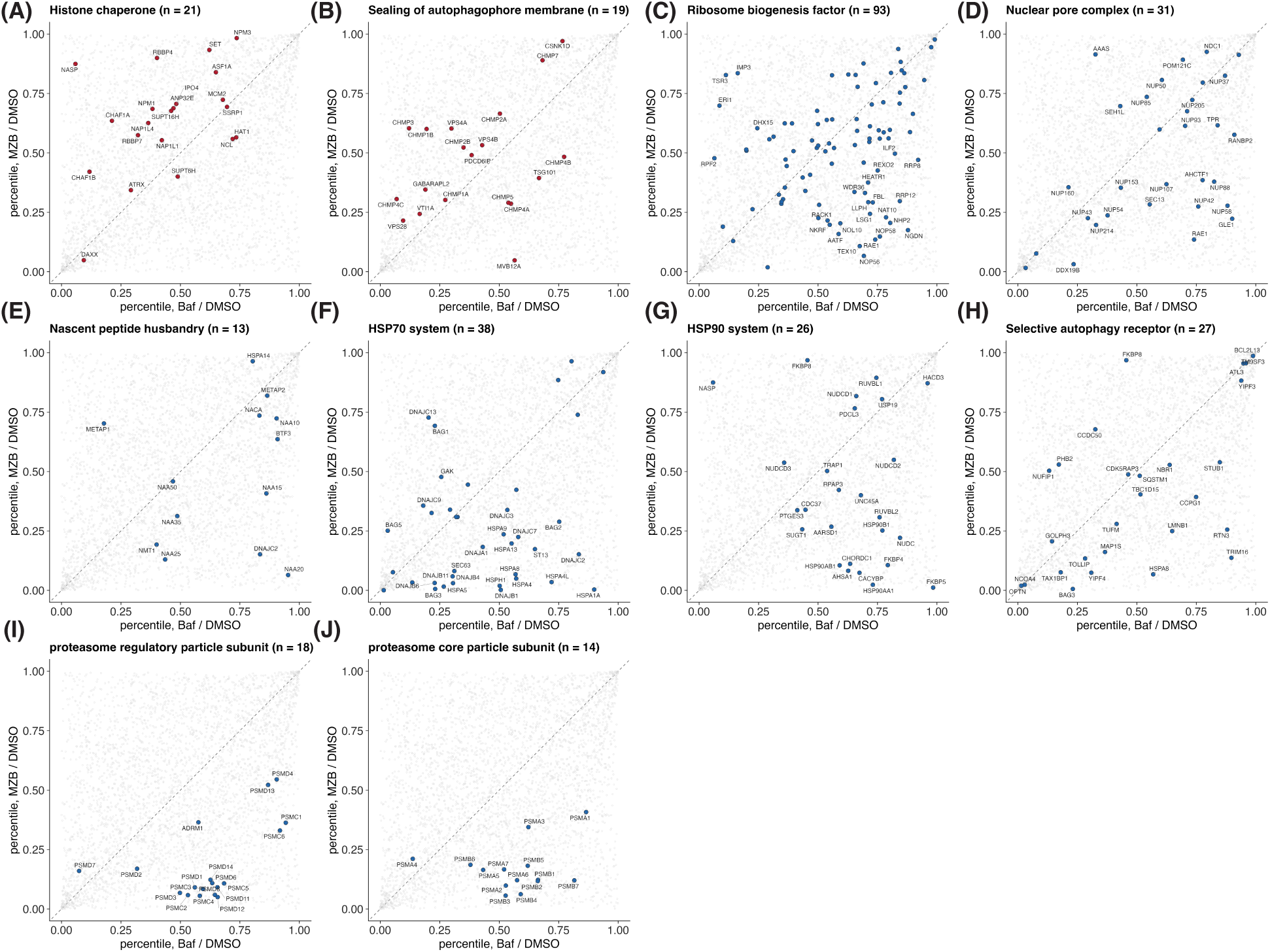
Degradation preference maps for individual proteostasis network groups. Percentile responses under MZB and BafA1 are shown for members of each PN group (A to J) overlaid on the full quantified proteome in grey. The group name and number of quantified members are given in each panel title. Point color indicates the direction of the group level sign test, with blue indicating ALP preference and red indicating UPS preference. The dashed line is the identity diagonal, and individual proteins are labeled with gene symbols.

## Reference

(1) Powers, E. T.; Morimoto, R. I.; Dillin, A.; Kelly, J. W.; Balch, W. E. Biological and chemical approaches to diseases of proteostasis deficiency. Annu Rev Biochem 2009, 959–991.

(2) Labbadia, J.; Morimoto, R. I. The biology of proteostasis in aging and disease. Annu Rev Biochem 2015, 435–464.

(3) Klaips, C. L.; Jayaraj, G. G.; Hartl, F. U. Pathways of cellular proteostasis in aging and disease. J Cell Biol 2018, 51–63.

(4) Hershko, A.; Ciechanover, A. The ubiquitin system. Annu Rev Biochem 1998, 425–479.

(5) Mizushima, N.; Komatsu, M. Autophagy: renovation of cells and tissues. Cell 2011, 728–741.

(6) Kaushik, S.; Cuervo, A. M. The coming of age of chaperone-mediated autophagy. Nat Rev Mol Cell Biol 2018, 365–381.

(7) Boisvert, F. M.; Ahmad, Y.; Gierlinski, M.; Charriere, F.; Lamont, D.; Scott, M.; Barton, G.; Lamond, A. I. A quantitative spatial proteomics analysis of proteome turnover in human cells. Mol Cell Proteomics 2012, M111 011429.

(8) Mathieson, T.; Franken, H.; Kosinski, J.; Kurzawa, N.; Zinn, N.; Sweetman, G.; Poeckel, D.; Ratnu, V. S.; Schramm, M.; Becher, I.;, et al. Systematic analysis of protein turnover in primary cells. Nat Commun 2018, 689.

(9) Potts, B. C.; Albitar, M. X.; Anderson, K. C.; Baritaki, S.; Berkers, C.; Bonavida, B.; Chandra, J.; Chauhan, D.; Cusack, J. C.; Fenical, W.;, et al. Marizomib, a proteasome inhibitor for all seasons: preclinical profile and a framework for clinical trials. Curr Cancer Drug Targets 2011, 254–284.

(10) Yoshimori, T.; Yamamoto, A.; Moriyama, Y.; Futai, M.; Tashiro, Y. Bafilomycin A1, a specific inhibitor of vacuolar-type H(+)-ATPase, inhibits acidification and protein degradation in lysosomes of cultured cells. J Biol Chem 1991, 17707–17712.

(11) Mauvezin, C.; Neufeld, T. P. Bafilomycin A1 disrupts autophagic flux by inhibiting both V-ATPase-dependent acidification and Ca-P60A/SERCA-dependent autophagosome-lysosome fusion. Autophagy 2015, 1437–1438.

(12) Salovska, B.; Li, W.; Bernhardt, O. M.; Germain, P. L.; Wang, Q.; Gandhi, T.; Reiter, L.; Liu, Y. A robust multiplex-DIA workflow profiles protein turnover regulations associated with cisplatin resistance and aneuploidy. Nat Commun 2025, 5034.

(13) Aniento, F.; Roche, E.; Cuervo, A. M.; Knecht, E. Uptake and degradation of glyceraldehyde-3-phosphate dehydrogenase by rat liver lysosomes. J Biol Chem 1993, 10463–10470.

(14) Dikic, I. Mechanisms controlling EGF receptor endocytosis and degradation. Biochem Soc Trans 2003, 1178–1181.

(15) Ding, W. X.; Li, M.; Biazik, J. M.; Morgan, D. G.; Guo, F.; Ni, H. M.; Goheen, M.; Eskelinen, E. L.; Yin, X. M. Electron microscopic analysis of a spherical mitochondrial structure. J Biol Chem 2012, 42373–42378.

(16) Wyant, G. A.; Abu-Remaileh, M.; Frenkel, E. M.; Laqtom, N. N.; Dharamdasani, V.; Lewis, C. A.; Chan, S. H.; Heinze, I.; Ori, A.; Sabatini, D. M. NUFIP1 is a ribosome receptor for starvation-induced ribophagy. Science 2018, 751–758.

(17) Pakos-Zebrucka, K.; Koryga, I.; Mnich, K.; Ljujic, M.; Samali, A.; Gorman, A. M. The integrated stress response. EMBO Rep 2016, 1374–1395.

(18) Costa-Mattioli, M.; Walter, P. The integrated stress response: From mechanism to disease. Science 2020.

(19) Elsasser, S.; Elia, L. P.; Morimoto, R. I.; Powers, E. T.; Finley, D.; Costa, B.; Budron, M.; Tokuno, Z.; Wang, S.; Iyer, R. G.;, et al. A Comprehensive Enumeration of the Human Proteostasis Network. 2. Components of the Autophagy-Lysosome Pathway. bioRxiv 2023.

(20) Toyama, B. H.; Savas, J. N.; Park, S. K.; Harris, M. S.; Ingolia, N. T.; Yates, J. R.; Hetzer, M. W. Identification of long-lived proteins reveals exceptional stability of essential cellular structures. Cell 2013, 971–982.

(21) Marshall, R. S.; Li, F.; Gemperline, D. C.; Book, A. J.; Vierstra, R. D. Autophagic Degradation of the 26S Proteasome Is Mediated by the Dual ATG8/Ubiquitin Receptor RPN10 in Arabidopsis. Mol Cell 2015, 1053–1066.

(22) Cohen-Kaplan, V.; Livneh, I.; Avni, N.; Fabre, B.; Ziv, T.; Kwon, Y. T.; Ciechanover, A. p62- and ubiquitin-dependent stress-induced autophagy of the mammalian 26S proteasome. Proc Natl Acad Sci U S A 2016, E7490–E7499.

(23) Marshall, R. S.; Vierstra, R. D. Dynamic Regulation of the 26S Proteasome: From Synthesis to Degradation. Front Mol Biosci 2019, 40.

(24) Thoreen, C. C.; Sabatini, D. M. Rapamycin inhibits mTORC1, but not completely. Autophagy 2009, 725–726.

(25) Claydon, A. J.; Beynon, R. Proteome dynamics: revisiting turnover with a global perspective. Mol Cell Proteomics 2012, 1551–1565.

(26) Ross, A. B.; Langer, J. D.; Jovanovic, M. Proteome Turnover in the Spotlight: Approaches, Applications, and Perspectives. Mol Cell Proteomics 2021, 100016.

(27) Ma, Y.; Yates, J. R., 3rd. Proteomics and pulse azidohomoalanine labeling of newly synthesized proteins: what are the potential applications? Expert Rev Proteomics 2018, 545–554.

(28) Kirschner, F.; Arnold-Schild, D.; Leps, C.; Lacki, M. K.; Klein, M.; Chen, Y.; Ludt, A.; Marini, F.; Kucuk, C.; Stein, L.;, et al. Modulation of cellular transcriptome and proteome composition by azidohomoalanine-implications on click chemistry-based secretome analysis. J Mol Med (Berl) 2023, 855–867.

(29) Békés, M.; Langley, D. R.; Crews, C. M. PROTAC targeted protein degraders: the past is prologue. Nat Rev Drug Discov 2022, 181–200.

(30) Ji, C. H.; Kim, H. Y.; Lee, M. J.; Heo, A. J.; Park, D. Y.; Lim, S.; Shin, S.; Ganipisetti, S.; Yang, W. S.; Jung, C. A.;, et al. The AUTOTAC chemical biology platform for targeted protein degradation via the autophagy-lysosome system. Nat Commun 2022, 904.

